# Effect Of Inflammatory Pain On Alcohol-Induced Dopamine Release In The Nucleus Accumbens: Behavioural Implications In Rat Models

**DOI:** 10.1101/862318

**Authors:** Yolanda Campos-Jurado, Jesús David Lorente, José Luis González-Romero, Luis Granero, Ana Polache, Lucía Hipólito

**Affiliations:** Department of Pharmacy and Pharmaceutical Tech. and Parasit. University of València, Spain

## Abstract

Recent studies have drawn the attention to the link between Alcohol Use Disorder (AUD) and the presence of pain. Indeed, the correct management of pain in patients with a previous history of AUD has been reported to decrease the risk of relapse in alcohol drinking, suggesting that in this prone population, pain may increase the vulnerability to relapse. Previous data in male rats revealed that inflammatory pain desensitizes mu opioid receptors (MORs) in the ventral tegmental area (VTA) and increases intake of high doses of heroine. Due to the relevant role of MORs in alcohol effects, we hypothesize that pain may also alter alcohol reinforcing properties and therefore affect alcohol relapse in male rats. Our microdialysis studies show that the presence of inflammatory pain blunted the increase of extracellular dopamine levels in the Nucleus Accumbens induced by 1.5g/kg of ethanol (s.c.). Moreover, we also revealed that the administration of 52 nmol of ethanol into the VTA failed to induce place preference only in inflammatory pain-suffering animals, and a higher dose (70nmol) was necessary to reverse this effect. Finally, we evaluated the effect of inflammatory pain on the alcohol deprivation effect (ADE) in long-term ethanol-experienced male rats. After four cycles of free ethanol intake and abstinence periods, inflammatory pain induced ADE without affecting its magnitude. These intriguing data reveals the impact of pain on neurochemical and behavioral effects following alcohol administration but also underscore the necessity of finding an appropriate paradigm to determine the long-term behavioral consequences.

## 1. Introduction

Chronic pain is a significant contributor to the worldwide burden of disease, affecting up to 30% of the population in the United States [35]. Recent epidemiological studies reveal a possible link between the presence of pain and Alcohol Use Disorder (AUD) [1]. Patients with a history of AUD usually suffer from pain syndromes during withdrawal, being painful peripheral neuropathy one of the most common complications [7,9]. Conversely, there is also evidence that the presence of pain may alter alcohol use patterns. In fact, treatment-seeking alcoholics have a highly prevalence of pain compared to the rest of the population [2,36] and higher levels of pain correlate with higher risk of alcohol relapse [15]. This interplay between pain and addiction has been commonly reported for opioids [5,21] and it has also been thoroughly investigated in preclinical studies.

There is evidence that pain causes alterations in the mesocorticolimbic system (MCLS), a key region in the modulation of affection, motivation and drug addiction [19,32]. In particular, inflammatory pain desensitizes mu opioid receptors (MORs) in the MCLS, what reduces morphine-induced conditioned place preference (CPP) at regular doses [22,25] and drives to the transition into a higher heroin dose consumption [14]. Many similarities between opioids and ethanol mechanism of action have been profoundly evidenced and it is well established that MORs play a critical role in ethanol action over the MCLS (see Nutt 2014 for review [23]). Indeed, different behavioral approaches have described that blockade of local MORs in the ventral tegmental area (VTA) reduces or suppresses ethanol-induced reinforcing behaviors [11,17,27]. Therefore, if the presence of pain induces alterations in the function of the MORs located in the MCLS it could be possible that it also affects ethanol action over this dopaminergic pathway. As a consequence, pain may also, in some way, have an effect on ethanol reinforcing properties and relapse. To this extent, some preclinical research has recently shown an increase in voluntary ethanol consumption in pain-suffering male mice when compared to pain-free groups [4,39].

However, there is still a lack of preclinical data exploring the possible effect of pain on ethanol relapse related behaviors. Relapse constitutes a major obstacle for AUDs treatments, hence at most 50% of treated people do not achieve remission after long follow up periods [10]. Moreover, and as previously mentioned, clinical studies have shown that the correct management of the pain situation resulting in decreases in pain levels are associated with lower risk of alcohol relapse [15]. Therefore, it is worth to further investigate whether pain condition can constitute a risk factor to ethanol relapse and if so, which could be the contributing neural mechanisms. Although different behavior may be observed depending on the sex, based in previous data that have shown an effect in alcohol intake in males but not in females, we decided to perform our studies in male rats [39]. To this end, in the present study we evaluate inflammatory pain-induced changes in: 1) ethanol-evoked dopamine (DA) release in the nucleus accumbens (NAc); 2) context-induced associations by analyzing intra-VTA ethanol induced place preference; and 3) ethanol-relapse by evaluating the appearance of the Alcohol Deprivation Effect (ADE) in long-term ethanol experienced rats.

## 2. Methods

### I. Rats housing

Male albino Wistar rats (Envigo, Spain. 300-340 g at the time of surgery or 280-300 g at the beginning of the long-term ethanol non-operant self-administration) were individually housed in plastic cages (48 × 38 × 21 cm^3^) provided with shredded aspen bedding (Teklad) and cotton enrichment (iso-BLOX^™^, Teklad) and controlled humidity and temperature (22°C), a 12:12-h light/dark cycle (on 08:00, off 20:00), and free access to food (Teklad Global Diets) and tap water. All the procedures were carried out in strict accordance with the EEC Council Directive 86/609, Spanish laws (RD 53/2013) and animal protection policies. Experiments were approved by the Animal Care Committee of the University of Valencia and authorized by the Regional Government.

### II. Surgery and inflammatory pain model

All surgeries were performed under isoflurane (1.5 MAC) anesthesia under aseptic conditions. For microdialysis experiments, rats were implanted stereotaxically (Stoelting, USA) with bilateral vertical concentric-style microdialysis probes containing 2 mm of active membrane (Hospal AN69; molecular cutoff 60,000 Da) constructed according to Santiago and Westerink 1990 [29] into the NAc (anteroposterior: +1.5 mm, mediolateral: ±1.6 mm and dorsoventral = −8.0 mm from bregma) [26]. For CPP experiments rats were implanted bilaterally with two 28-gauge guide cannulae aimed at 1.0 mm above the posterior VTA (anteroposterior: −6.0 mm, mediolateral: ±1.8 mm and dorsoventral: −7.9 mm). Both coordinates have been successfully used in previous studies in our laboratory [12,13]. Cannulae were angled toward the midline at 10° from the vertical. A stainless-steel stylet (33-gauge), extending 1.0 mm beyond the tip of the guide cannula, was put in place at the time of surgery and removed at the time of testing.

We selected the Complete Freund Adjuvant (CFA) model of inflammatory pain. CFA (Calbiochem) was diluted in the same volume of sterile saline before its subcutaneous injection of 0.1 ml in the plantar surface of the hindpaw [14]. The injection of saline or CFA in the hindpaw was performed at the same time of the surgery for the microdialysis experiments, four days after the cannulation surgery for the CPP experiments and two days before the re-introduction of ethanol for the ADE experiment.

### III. Microdialysis

Forty-eight hours after the sterotaxical surgery animals belonging both to the pain (n=9) and pain-free (n=8) group were placed in Plexiglas bowls. A PE10 inlet tubing was attached to a 2.5 mL syringe (Hamilton) mounted on a syringe pump (Harvard instruments, South Natick, MA, USA) and connected to the probes. Probes were continuously perfused with artificial cerebrospinal fluid (aCSF) comprising 0.1 mM aqueous phosphate buffer containing 147 mM NaCl, 3.0 mM KCl, 1.3 mM CaCl2 and 1.0 mM MgCl2 (pH 7.4) at a flow rate of 3.5 μL/min. Following a minimum stabilization period of 1 hour, samples were collected every 20 min and extracellular DA levels were determined by using offline HPLC with electrochemical detection. The HPLC system consisted of a Waters 510 series pump in conjunction with an electrochemical detector (Mod. Decade, Antec, Leyden, The Netherlands). The applied potential was = 0.55 V (ISAAC cell, Antec, Leyden, The Netherlands). Dialysates were injected into a 2.1 mm RP-18 column (Phenomenex, Gemini-NX 3 µm 100 × 2.00 mm) with a 65 μL sample loop. The mobile phase consisted of a sodium acetate/acetic acid buffer (0.05 mol/L, pH = 6) containing 140 mmol/L of sodium chloride, 200 mg/L of 1-octanesulfonic acid, 100 mg/L of EDTA and 150 mL/L of methanol. The mobile phase was pumped through the column at a flow rate of 0.06 mL/min. Chromatograms were analyzed and compared with standards (1.1, 2.2, 5.5 and 11 nM) run separately on each experimental day, using the AZUR 4.2 software (Datalys, France). Once DA baseline level was established (defined as 3 consecutive samples with <10% variation in DA content), 2 mL of saline were subcutaneously (s.c.) administered and DA levels were analyzed every 20 min for 100 min as a control for the possible effect of the s.c. injection itself. Right after, a single s.c. ethanol dose (1.5 g/kg diluted in 2mL of saline) was administered and DA levels in the dialysates were analyzed for 160 min more.

At the end of the microdialysis experiments, rats were sacrificed and the brains removed and rapidly frozen in dry ice; 40-μm-thick coronal slices of the NAc core were obtained using a cryostat and stained with a cresyl violet protocol to verify proper probe placement.

### IV. Ethanol Conditioned Place Preference

The CPP test was performed in a non-biased two-compartment box connected by a removal barrier with an open door in the middle. The two compartments differed by the wall color: black and white vertical stripes (vertical compartment) and black and white horizontal stripes (horizontal compartment). Four days after recovery from surgery and two days prior to the “Pretest”, rats received a subcutaneous administration in the hindpaw of 0.1 mL saline or 0.1 mL CFA. The next day animals were exposed to the CPP box for 5 min in order to habituate them to the apparatus. The day prior to the conditioning, animal natural preference for the compartments was tested during 15 min (Pretest). During “Conditioning”, rats received bilateral intra-VTA infusions before placing them in the appropriate compartment. All conditioning phases consisted on 8 sessions (2 sessions/day: morning session and afternoon session) of 30 min distributed in 4 days. 27 animals were randomly assigned to one of the ethanol doses tested (70 or 52 nmol) and the exposure to conditioning compartments was counterbalanced in both groups. Control group animals (n=6) received 8 administrations of the equivalent volume of aCSF meanwhile the other groups received ethanol or aCSF on alternate sessions. After the last conditioning session, each animal was subjected to the “Test” for its place preference: the rat was placed in the open door of the barrier, and the time spent in each compartment was recorded over 15 min. Place preference scores were calculated as test minus pretest time spent (in seconds) on the ethanol-paired compartment.

At the end of the CPP experiments, rats were sacrificed and the brains removed and rapidly frozen in dry ice; 40-μm-thick coronal slices of the VTA were obtained using a cryostat and stained with a cresyl violet protocol to verify proper cannulae placement. This CPP procedure is summarized in Figure 2A.

### V. Long-term ethanol non-operant self-administration

Before initiating the long-term voluntary ethanol drinking procedure, rats (n=20) were habituated to the animal room for two weeks. Next, animals were given continuous access to tap water and to 5 %, 10 %, and 20 % (v/v) ethanol solutions in their home cages. Every week animals were weighted; all drinking solutions renewed and changed the positions of the four bottles to avoid location preferences. After 8 weeks of continuous ethanol availability, the first 2-week deprivation period was introduced. Following, rats were given access to alcohol again and three more deprivation periods were performed in a random manner. The duration of these drinking and deprivation periods was irregular: 6 ± 2 weeks and 2 ± 1 weeks, respectively, in order to prevent behavioral adaptations [24]. The total amount of ethanol intake (g/Kg/day) was recorded during the whole experiment by weighing the bottles. Animals were randomly assigned to one of the two experimental groups. The baseline drinking for each group was considered as the average of the measurements of the three last days prior to the 4th abstinence period. During this last abstinence period and 48h before re-introduction of the ethanol solutions, rats received a subcutaneous administration in the hindpaw of 0.1 ml saline or 0.1 mL CFA. After the re-introduction of the ethanol solutions, the daily weighing routine was restored during the three post-abstinence days in order to assess the ADE.

### VI. Supervision of the Inflammatory pain model

With the objective to assess the level of inflammation induced by CFA injection we measured the dorso-ventral distance of the rats injected hindpaw and compared it to the contralateral hindpaw distance. This measurement was performed right before sacrifice in both, the microdialysis and the long-term non-operant self-administration experiments.

For the CPP experiment, we tested the mechanical nociception thresholds by using the Von Frey test before and after 2 and 9 days of intraplantar injections (after the pretest and test sessions). Following 20 min of habituation to the apparatus, nociception thresholds were measured by the manual application of five filaments (Aesthesio®) with a simplified up-down method, as described in Bonin 2014 [3]. Results were expressed by the mean of nociception threshold (in grams, g).

### VII. Statistical methods

All data sets are expressed as Mean ±SEM. In all cases homogeneity of variance was tested and the significance level was always set at p < 0.05.

#### Microdialysis experiments

DA levels in the three dialysate samples defining the baseline conditions (expressed as fmol in 65 μl), were averaged to calculate baseline levels in each animal. Differences in baseline levels between groups (saline and CFA treated animals) were evaluated using the unpaired Student’s *t* test.

DA levels were transformed to percentages of baseline for each individual rat and were statistically analyzed by a mixed two-way ANOVA for repeated measures, with group (saline or CFA injection) taken as the between-subjects factor and time as the within-subjects factor. Significant interaction time × group was further analyzed by means of a Bonferroni correction for multiple comparisons. Significant effects of time were analyzed by one-way ANOVA with repeated measures per each group followed by Bonferroni multiple-comparisons test and post-injection values were compared to the last baseline measure (t=0 in Figure 1B).

**Figure 1.**
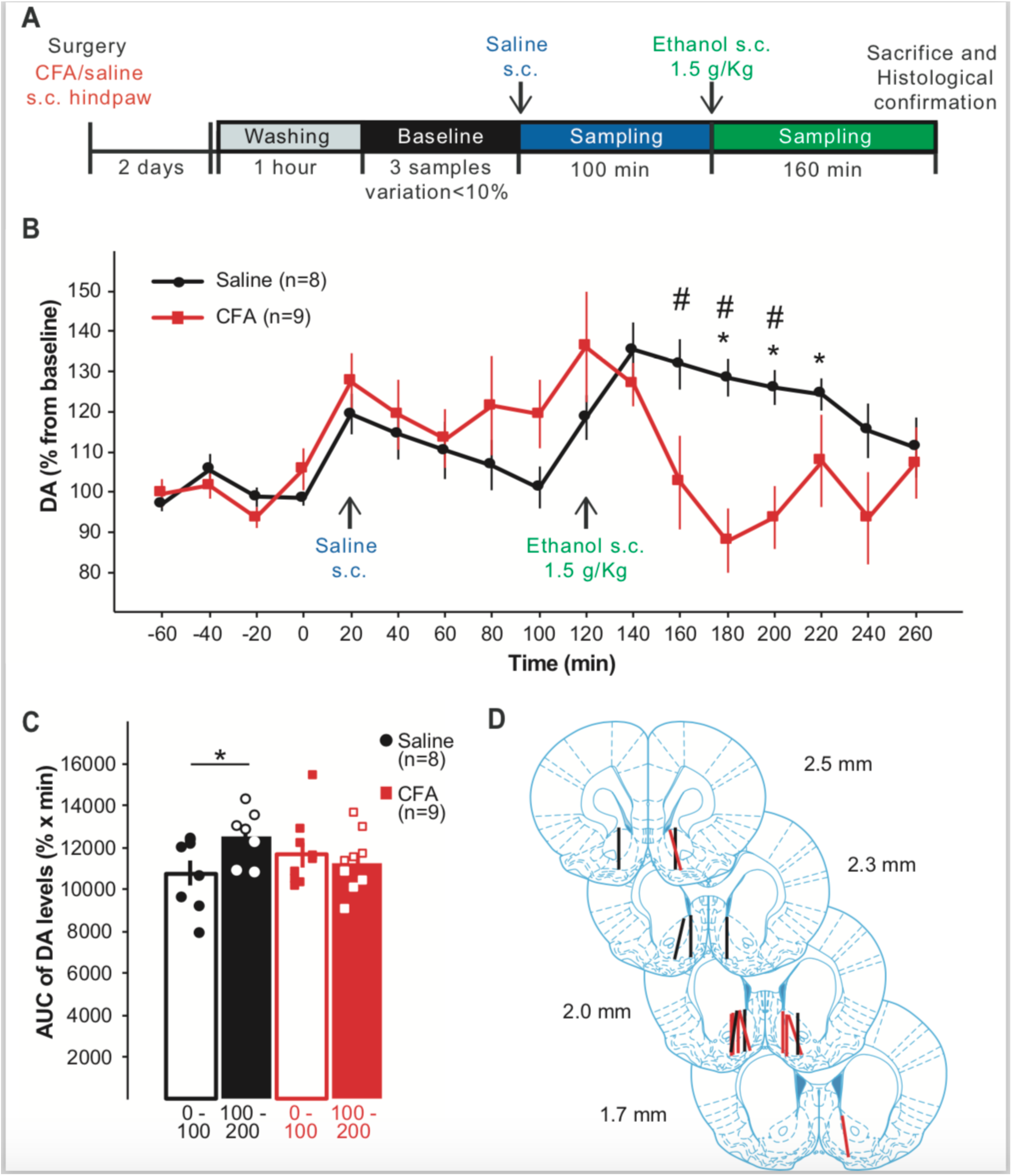
Pain decreases ethanol evoked DA release in the NAc. **A.** Schematic of the experimental design. **B.** DA extracellular levels in the NAc. Data are mean and SEM represented as percentage from baseline; # denotes significant differences between groups (Bonferroni multiple comparison, p < 0.05). * denotes significant differences in the within-subjects effect of time (Bonferroni multiple comparisons, p < 0.05). **C.** Global change in DA levels induced by saline (0 −100 min) and ethanol (100-200 min) calculated as AUC. * denotes significant differences (two-way ANOVA for repeated measures followed by Bonferroni multiple comparisons, p<0.05). **D.** Diagram of coronal sections indicating the placement of microdialysis probes in the NAc for control (black) and CFA-treated (red) rats in the NAc. Bars could represent more than one or probe placement.

**Figure 2.**
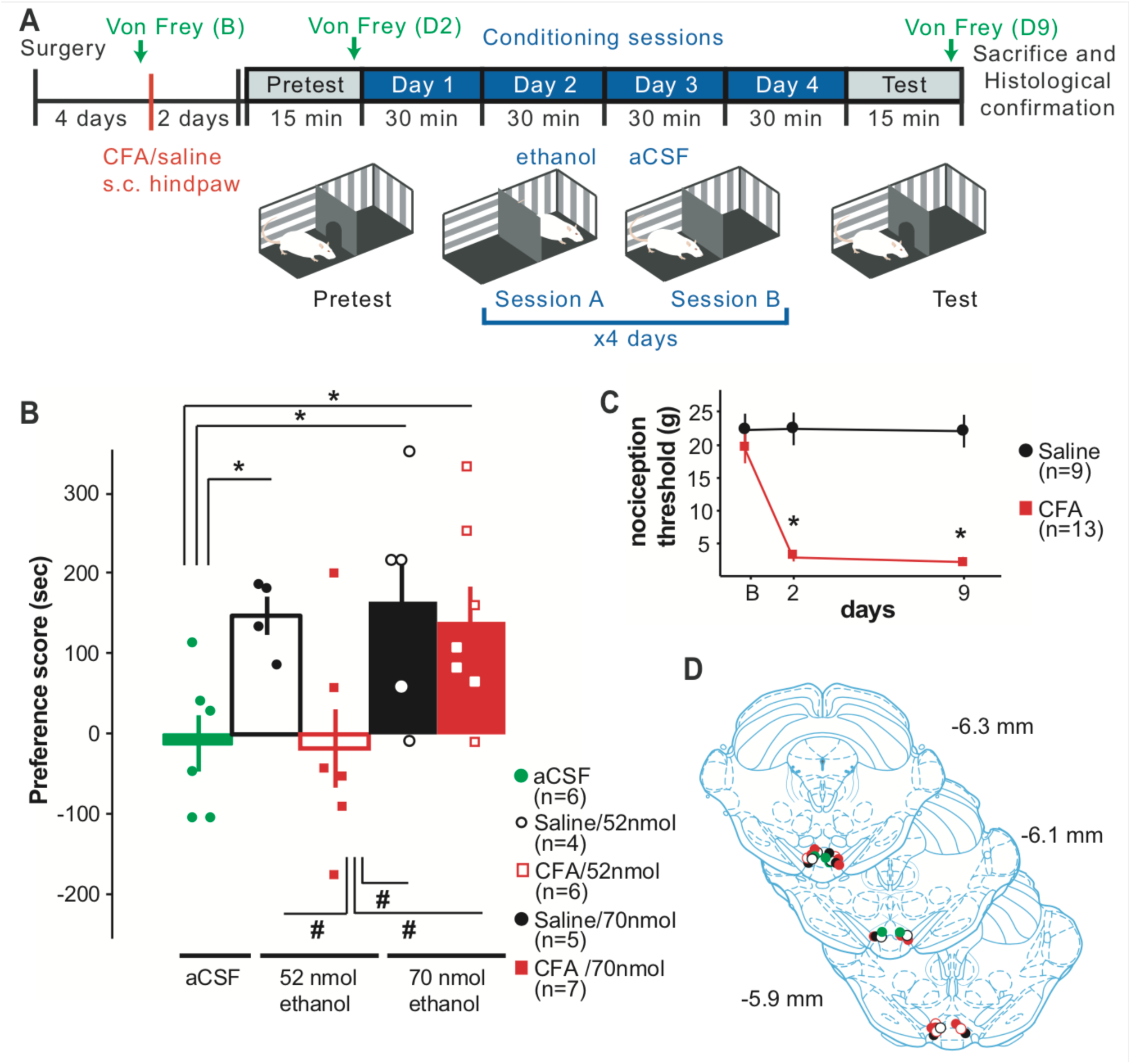
Pain alters intra-VTA ethanol induced CPP. **A.** Schematic of the experimental design. **B.** Preference score induced by the intra-pVTA administration of aCSF (green) or ethanol (52 and 70 nmol) in saline (black) and CFA (red) treated rats. Data are mean and SEM and represent preference scores, calculated as test minus pretest time spent on the ethanol pairedcompartment (n= 5-7/group). * denotes significant differences from control group (aCSF) and # denotes significant differences from CFA-treated rats receiving 52 nmol of ethanol (one-way ANOVA followed by LSD multiple comparisons). **C.** Mean and SEM of the paw withdrawal thresholds measurements in black for the saline-treated group and in red for the CFA-treated group. * denotes significant differences between groups and # denotes significant differences from baseline (two-way ANOVA for repeated measurements followed by Bonferroni multiple comparisons, p<0.001). **D.** Diagram of coronal sections indicating microinjection cannula tips in VTA for control (green), saline (black) and CFA-treated (red) rats in the intra-VTA injection. Dots could represent more than one or cannula placement.

Areas under the curve (AUC) were calculated from 0 to 100 min (post-saline effect) and from 100 to 200 min (post-ethanol effect) for each rat from percentage data. AUCs were statistically analyzed by mixed two-way ANOVA with repeated measures with group as a between-subjects factor and treatment (saline and ethanol) as a within-subjects factor followed by Bonferroni corrections for multiple comparisons when interactions were found to be significant.

#### Conditioned place preference

For CPP experiments results are expressed in preference score, calculated as time spent in ethanol paired compartment during test minus time spent in the same compartment during pretest. Preference scores were analyzed using one-way ANOVA, followed by LSD test.

For the Von Frey test results are expressed as mechanical nociceptive threshold in grams. Nociceptive thresholds were statistically analyzed by mixed two-way ANOVA with repeated measures with group (CFA or saline) as a between-subjects factor and time as a within-subjects factor followed by Bonferroni corrections for multiple comparisons.

#### Ethanol non-operant self-administration

Data were expressed as total amount of ethanol intake (g/Kg/day). Basal and either combined and individual measures of post-abstinence ethanol intake were statistically analyzed by mixed two-way ANOVA with repeated measures with group as a between-subjects factor and period (basal and post-abstinence) as a within-subjects factor followed by Bonferroni corrections for multiple comparisons when interactions were found to be significant.

## 3. Results

### 3.1 Effect of inflammatory pain on ethanol-evoked DA release in the NAc

The objective of this experiment was to analyze whether inflammatory pain affects ethanol-induced DA signaling within the NAc. For that, forty-eight hours after CFA injection extracellular DA content in the accumbal dialysates was monitored every 20 min. Once a stable baseline for DA was achieved (<10% variation in three consecutive samples), rats received a first saline s.c. injection to discriminate the possible effect of the manipulation/injection itself on the NAc DA levels. Next, a single ethanol dose was administered (1.5 g/kg, s.c.) and DA levels were monitored until the end of the experiment for a total of 260 min. Baseline DA levels (mean ± SEM) were 51.8 ± 9.8 fmol/65 μl and 62.8 ± 8.4 fmol/65 μl for the control versus pain group, respectively. Our results indicate that the presence of inflammatory pain does not have a significant effect on basal extracellular DA levels (*t* test, *p*=0.517 n=8-9).

The mixed ANOVA for repeated measures detected a significant effect in the within-subject variable. Indeed, both the time (*F*_(16,240)_= 4.195, *p*<0.001), in the interaction time × group (*F*_(16,240)_=3.506, *p*<0.001), but not in the effect of group (*F*_(1,15)_=1.140, p=0.302). The analysis of the effect in the within-subject variable time was further analyzed by means of a one-way ANOVA for repeated measures. As expected, in pain free animals 1.5 g/kg ethanol dose induced a significant increase in DA release up to 135% from 80 min to 120 min (180 to 220 min time points in 1B) after its administration (one-way ANOVA for repeated measures, within-subjects effect of time *F*_(16,112)_=7.631, *p*<0.001; Bonferroni multiple comparisons versus the last basal time point, t=0 min: *p*_180_=0.023, *p*_200_=0.017, *p*_220_=0.015). Interestingly, although CFA-treated rats experienced an ethanol-induced DA release (136% from baseline), this increase was not significantly higher compared to last value of baseline (one-way ANOVA for repeated measures, within-subjects effect of time *F*_(16,128)_=3.081 p= 0.0001, Bonferroni multiple comparisons versus t=0 min: *p*<0.001, *p*_20_-*p*_260_>0.05). Indeed, ethanol-evoked DA levels in the CFA-treated group were significantly lower than the saline group at 60, 80 and 100 min after 1.5 g/kg ethanol administration (Figure 1B; two-way ANOVA for repeated measures, interaction time × group *F*_(16,240)_=3.506, *p*<0.001; Bonferroni correction for multiple comparisons, *p*_60_=0.048, *p*_80_=0.001, *p*_100_=0.002). Finally, it is important to mention that Saline s.c. injection induced a slight increase up to 127% and 119% in DA release (at 20 min time point) both in pain and pain-free animals, respectively, although this increase in DA levels was no statistically significant (pain group: *p*_20_=0.688, pain-free group: *p*_20_=0.896; compared to last baseline point t=0).

To further quantify this inflammatory pain-induced effect on the ethanol-evoked DA release, we calculated the AUC for the following 100 min after each administration in both pain and pain-free groups (Figure 1C). This parameter allowed us to compare the global response to ethanol on extracellular NAc DA levels. Mean AUC values were significantly different between saline and CFA-treated groups (two-way ANOVA for repeated measures, within-subjects interaction time × group *F*_(1,15)_= 6.384, *p=*0.023). The ethanol-induced total effect in the pain-free group was significantly higher than the total effect induced by the saline injection (Bonferroni correction for multiple comparisons, *p*=0.013), whereas no significant differences were found between saline and ethanol treatments in pain animals (Bonferroni correction for multiple comparisons, *p*=0.502).

Only animals that showed correct placement of the microdialysis probe in the NAc are included in the analysis. Position of the active portion of the dialysis membrane can be inspected in Figure 1D.

### 3.2 Effect of inflammatory pain on ethanol-induced CPP

In this experiment, we used the CPP paradigm to assess whether the above reported pain-induced changes in ethanol-evoked DA release could be reflected also in changes on ethanol-induced context learned associations. For that, we analyzed the ability of two different ethanol doses directly administered into the posterior VTA to induce CPP in animals under pain and pain-free condition. Our results show that inflammatory pain causes significant alterations on ethanol-induced CPP (Figure 2B, one-way ANOVA *F*_(4,23)_=3.685, *p*=0.018). Concretely, pain-free animals that received the lowest ethanol dose (52 nmol) showed a preference score significantly higher than the control group (146 ± 26 vs −12 ± 36, LSD test, p=0.040), whereas the group under inflammatory pain did not develop a preference for the ethanol paired compartment when compared to control group (−18 ± 49 vs −12 ± 36, LSD correction for multiple comparisons, p=0.920). Interestingly, the higher ethanol dose used (70 nmol) induced CPP in both saline and CFA-treated rats, shown as preference scores significantly higher than the control group (saline: 165 ± 54, p=0.016; CFA: 140 ± 44, p=0.024; LSD test). Moreover, the statistical analysis showed significant difference between doses only in CFA-treated group (CFA: p=0.019, saline: p=0.804).

Additionally, the results of the Von Frey test performed throughout the CPP experiment are represented in Figure 2C and confirmed that saline treated in the hind-paw animals maintained the mechanical nociceptive threshold as the one measured in the baseline session. On the contrary, animals that received CFA in the hindpaw showed a significant decrease in the mechanical nociceptive threshold that was maintained until the performance of the CPP test session (two-way ANOVA for repeated measures, within-subjects interaction time × group *F*_(2,44)_=16.299, *p*<0.001; Bonferroni correction for multiple comparisons, differences between groups: *pD2*<0.001 and *pD9*<0.001, differences from baseline, saline: *pD2*=1.000 and *pD9*=1.000, CFA: *pD2*<0.001 and *pD9*<0.001).

Only animals that showed correct placement of the cannula in the VTA were included in the analysis. Position of the cannulae tip can be inspected in Figure 2D.

### 3.3 Effect of inflammatory pain on ADE

The effect of inflammatory pain on ethanol relapse was studied by selecting a non-operant self-administration paradigm in which periods of continuous access to 4 different bottles (water and 5%, 10% and 20% (v/v) ethanol) were alternate with forced abstinence periods (access to water) (Figure 3A). During the last abstinence period and forty-eight hours before ethanol reintroduction, animals were injected with saline or CFA in the hindpaw and ADE was analyzed by measuring the average of ethanol consumption during the three days after reintroduction (Figure 3A). After the reintroduction of alcohol solutions both pain and pain-free groups showed an increase in alcohol consumption compared to basal (calculated as average of the last 3 days before the abstinence), indicating the occurrence of ADE and no significant differences were found between groups (two-way ANOVA for repeated measures, within subjects effect of time *F*_(1,18)_=45.599, *p*<0.001; interaction time × group *F*_(1,18)_=0.536, *p*=0.474). Therefore, Bonferroni correction for multiple comparisons showed significant difference between basal and post-abstinence ethanol consumption for both groups (*p*_saline_<0.001, *p*_CFA_=0.001). We also analyzed the individual ethanol intake for each of the three post-abstinence days (Figure 3B) and there was an increase after ethanol reintroduction in pain and pain-free groups, with no significant differences between groups (two-way ANOVA for repeated measures, within subjects effect of time *F*_(3,54)_=15.268, *p*<0.001; interaction time × group *F*_(3,54)_=0.306, *p*=0.821). Concretely, in pain-free animals, total ethanol intake was significantly higher in the 3 post-abstinence days compared to baseline (Bonferroni correction for multiple comparisons, *p*_day1_=0.001, *p*_day2_=0.019, *p*_day3_=0.026), whereas in pain group there were only significant differences between total ethanol intake in the first post-abstinence day compared to baseline (Bonferroni correction for multiple comparisons, *p*_day1_=0.001, *p*_day2_=0.061, *p*_day3_=1.000) (see Figure 3C).

**Figure 3.**
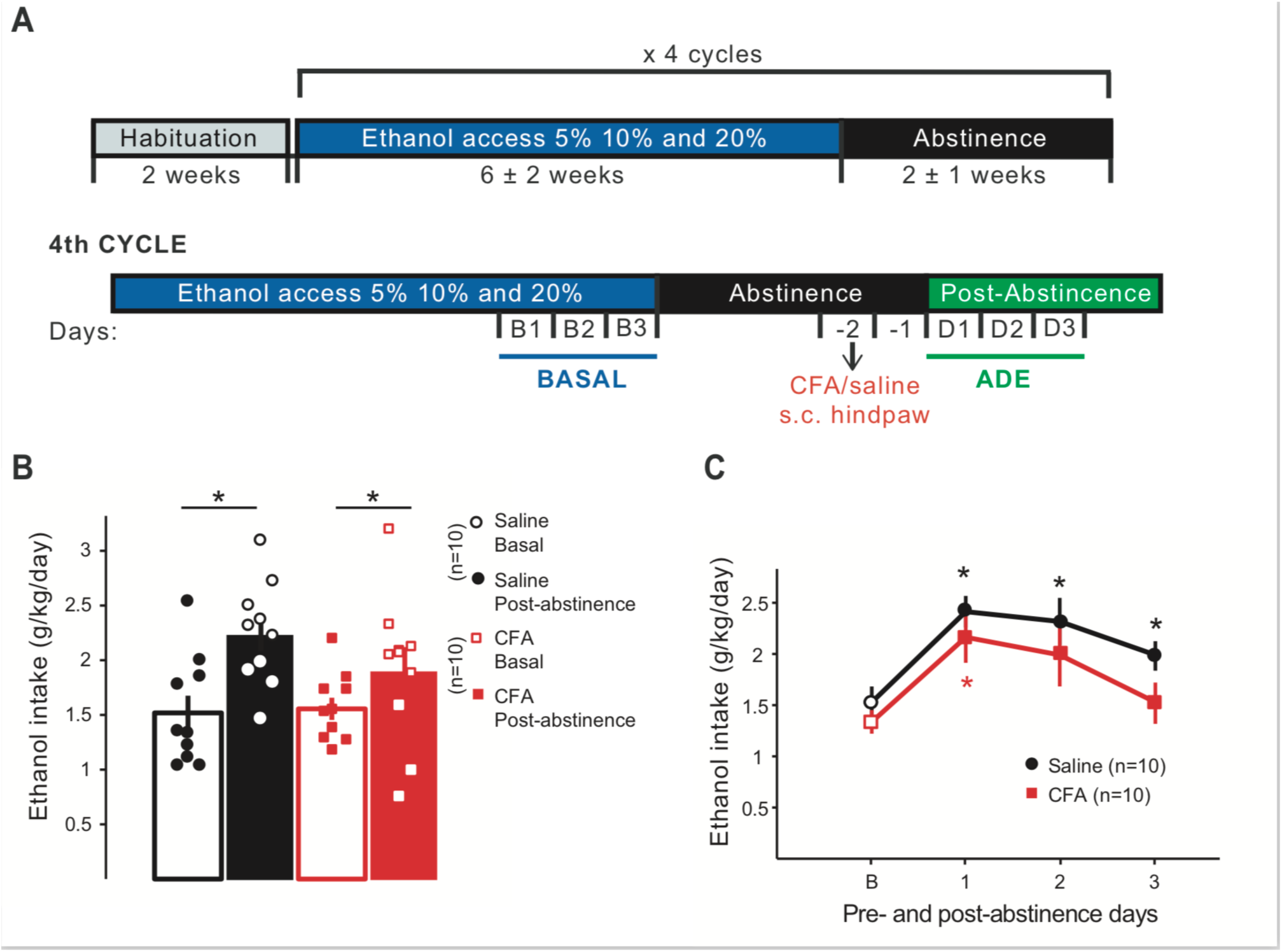
Inflammatory pain does not modify the magnitude of the Alcohol Deprivation Effect (ADE). **A.** Schematic of the experimental design. **B.** Mean and SEM of total alcohol intake (g/Kg/day) of the 3 days pre-(basal, empty bar) and post-abstinence (filled bars) shown in black for the saline-treated group (n= 10) and in red for the CFA-treated group (n=10). The two-way ANOVA for repeated measures showed a within subject effect of time (p<0.001). * denotes significant difference between basal and post-abstinence ethanol consumption (Bonferroni correction for multiple comparisons, p<0.001). **C.** Mean and SEM of total alcohol intake (g/kg/day) for basal and for each of the three post-abstinence days shown in black bar for the saline-treated group (n= 10) and in red bar for the CFA-treated group (n=10). The two-way ANOVA for repeated measures showed a within subject effect of time (p<0.001). * denotes significant differences from baseline (Bonferroni correction for multiple comparisons, p<0.05).

## 4. Discussion

In this paper, we present evidence supporting that both, ethanol-evoked neurochemical responses and dopamine-dependent behaviors, such as CPP, are altered in male pain-suffering animals, so that higher ethanol doses are needed to elicit similar effects to those observed in male pain-free rats. We also show that inflammatory pain, however, did not alter the size of ethanol relapse measured as the magnitude of ADE in long term ethanol-experienced male rats.

An association between pain condition, AUD and higher risk of relapse has been previously reported in the clinical literature [2,15,36]. In the preclinical setting, previous studies have showed that inflammatory pain is able to induce a loss of MORs function that results in alterations of these receptors regulation of the mesolimbic DA transmission. Inflammatory pain results in the intake of higher doses of heroin and promotes opioid dose escalation on a rat intravenous self-administration model [14]. Here, we provide further evidence on the effects that inflammatory pain exerts on DA mesolimbic responses evoked by drugs of abuse, in particular by another widely used drug: alcohol.

Pain and drug addiction share common neural circuits [1,9]. It has already been reported that brain pathways that mediate addiction and affection are altered by the presence of pain [5,18,21]. Opioid receptors, mainly MORs, located in the MCLS, are thought to mediate the reinforcing properties not only of opioids but also of ethanol, via regulation of DA extracellular levels within the NAc [6,37]. Previous research showed that heroin-evoked DA release in the NAc is blunted in rats under inflammatory pain condition, what drives to higher heroin dose consumption [14]. Similarly, in our experiment, the administration of ethanol did not provoke a significant increase in NAc DA levels in CFA-treated animals (Figures 1B and 1C). This finding further supports the fact that inflammatory pain alters the neurochemical response of the MCLS elicited by drugs. Curiously, the heroin-evoked increases of DA release described by Hipolito 2015 started 15 min after heroin administration and differences between pain-free and CFA-treated groups were found 30 and 45 min after this administration. According to our results, ethanol-evoked increases and differences between groups also appear later on time. In addition, the global effect elicited by ethanol in pain-free group was significantly higher than the saline-induced total effect. On the contrary, saline and ethanol-induced total effect were not statistically different in CFA-treated animals (Figure 1C), confirming that our results are derived from the drug pharmacological properties. Although the specific mechanism remains to be elucidated, our microdialysis results clearly show that inflammatory pain impairs ethanol-evoked DA release in the NAc, which may have an effect on the reinforcing properties of this drug.

It is classically accepted that drug-reward related behaviors (such as CPP) [30,38] are mediated by DA transmission within the NAc. Consequently, altered neurochemical function in MCLS can be translated into abnormal changes in these behaviors. Previous studies have reported that the presence of pain suppresses the preference for opioid-associated environments [22,25,31]. Again, our present results show that local administration of 52 nmol of ethanol intra-VTA is able to induce preference for the ethanol-paired compartment only in pain-free animals (Figure 2B). Very interestingly, we also show that a higher ethanol dose of 70 nmol is able to reverse this pain-induced blockade of ethanol place preference (Figure 2B), hence resulting in similar preference scores in both saline and CFA-treated groups. The shift of the behavior observed in CFA-treated rats depending on the intra-VTA dose of ethanol administered, provides more evidence that dopamine-dependent behaviors are altered in pain-suffering animals. In fact, in our experiments, 52 and 70 nmol of ethanol administered intra-VTA in pain-free animals shows a very similar behavioral output which is the induction of context-dependent associations. On the contrary, in pain animals the large difference observed in the behavioral output elicited by the two doses is showing a shift from non-active dose (52nmol) for these pain animals into an active dose (70 nmol). Moreover, our Von Frey test data confirms that no changes in the nociception where observed neither in the pain or nopain groups through the CPP experimental process, ruling out the possibility of unspecific behavioral effects derived from changes in mechanical nociception. These observed phenomena are, therefore, consistent with our previous findings reported in Hipolito et al. 2015. In that study, higher doses of heroin were necessary both to increase DA levels in the NAc and to elicit heroin self-administration in CFA-treated rats [14]. All in all, our results show that inflammatory pain alters ethanol reinforcing action on the MCLS which may have consequences in alcohol consumption patterns in male rats.

Lastly, we decided to study whether this previous reported alteration in ethanol-evoked neurochemical responses and dopamine-dependent behaviors induced by inflammatory pain could have an effect on AUDs related behaviors. Pain condition frequently elicits negative affective states driving to alterations in reward evaluation, decision making and motivation [1,34]. This pain induced-affective state has been reported to be mediated via the kappa opioid system [18]. AUD is also characterized by the abnormal persistence of negative affective states during withdrawal, that can promote drug seeking and relapse, probably through the dynorphin-kappa opioid system [8]. Thereupon, and given the recent epidemiological data showing that higher levels of pain correlate with a higher risk of alcohol relapse [15], we chose an alcohol relapse behavioral approach to further investigate this connection between pain and relapse. We selected a long-term non-operant self-administration paradigm that has been widely employed for the study of alcohol relapse-like behaviors in rodents by our and other groups and that provides predicted validity [20,24,33]. As expected, our results indicate that alcohol-deprived male rats under inflammatory pain developed ADE after alcohol reintroduction. The magnitude of the ADE did not change relative to pain-free animals. However, changes in the magnitude of the ADE are not representative of changes in alcohol relapse vulnerability. The ADE is defined as an increase in total ethanol intake that occurs during the first days after an abstinence period. It is true that numerous preclinical studies have used this paradigm to test different pharmacological strategies aimed to suppress or reduce ethanol relapse but, to our knowledge, it has never been used to show an increase of the risk of relapse [20,24]. Indeed, it is possible that pain does not induce an even higher ethanol intake but increases the vulnerability of relapse. In this case, in spite of differences in consumption, it would be plausible to expect higher rates of relapse in pain suffering animals. Therefore, further behavioral studies that allow to investigate alcohol drinking behavior in the face of vulnerability to relapse are needed and would shed more light in this aspect.

It also is important to note that pain-induced changes may differently affect addictive behaviors depending on the different stages of the AUD. In fact, previous studies using an intermittent two-bottle choice paradigm show that pain induction prior to alcohol exposure significantly increases total intake in male mice [4,39] and it may be plausible that pain increases alcohol intake during acquisition without modifying the magnitude of ADE. On the other hand, we have only tested male rats based in the previous data reported [4,39] and, it could be plausible that relapse-like behavior may be expressed differentially depending on the sex. In any case, our data highlight the necessity of finding the appropriate animal model that reflects the existing clinical evidence and allows us to study the alcohol-related behavioral implication of pain-induced alterations of the MCLS. Moreover, the present research provides relevant data as it analyses, for the first time, the effect of inflammatory pain on alcohol relapse in animal models and therefore constitutes an important contribution to the study of pain and AUD.

Altogether, our results reveal that the presence of inflammatory pain alters the response of the mesocorticolimbic dopaminergic pathway to alcohol. In fact, neurochemical and behavioral approaches point to the presence of inflammatory pain as a relevant factor in the neurobiological effects of alcohol and in alcohol addiction. The above-reported changes, however, did not allow us to model the observed increase vulnerability to relapse of pain patients in the clinical set up. These data further support the impact of pain on the reinforcing mechanisms following alcohol administration and also underscore the necessity and importance of finding an appropriate animal paradigm to determine the neural mechanisms underlying the drug-related behavioral consequences derived from pain.

## 5. Acknowledgments

This work was supported by Spanish Ministerio de Economia y Competitividad’’ MINECO PSI2016-77895-R (L.H.). We would like to thank Dr. Teodoro Zornoza for equipment and funding help. We also thank Ms. Pilar Laso for grant management and personnel of the Animal facilities at the University of Valencia for their help and effort in assuring animal welfare.

The authors declare no conflict of interest

## 6. Authors contribution

Conceptualization, L.H.; Methodology, Y.C-J., A.P., L.G., L.H; Formal Analysis, Y.C-J., A.P., L.G, L.H; Investigation, Y.C-J., J.D.L., J.L.G-R; Writing – Original Draft, Y.C-J, L.H.; Resources, L.H; Supervision, A.P., L.G., L.H. Writing – Review & Editing, Y.C.J., A.P., L.G., L.H.

